# Intronic miRNA Mediated Gene Expression Regulation Controls Protein Crowding Inside the Cell

**DOI:** 10.1101/286906

**Authors:** Prashant Sinha, Pragya Jaiswal, Ashwin K. Jainarayanan, Samir K. Brahmachari

## Abstract

**SUMMARY:** Gene regulatory effects of microRNAs at a posttranscriptional level has been established over the last decade. In this study, we analyze the interaction networks of mRNA translation regulation through intronic miRNA, under various tissue-specific cellular contexts, taking into account the thermodynamic affinity, kinetics, and the presence of competitive interactors. This database, and analysis has been made available through an open-access web-server, miRiam, to promote further exploration.

Here we report that expression of genes involved in Apoptosis Processes, Immune System Processes, Translation Regulator Activities, and Molecular Transport Activities within the cell are predominately regulated by miRNA mediation. Our findings further indicate that this regulatory effect has a profound effect in controlling protein crowding inside the cell. A miRNA mediated gene expression regulation serves as a temporal regulator, allowing the cellular machinery to temporarily ‘pause’ the translation of mRNA, indicating that the miRNA–mRNA interactions may be important for governing the optimal usage of cell volume.

## INTRODUCTION

The Human Genome Project has thrown up a surprise leading to discovery of many functional non-coding sequences in the human genome. Recent advances suggest that these non-coding sequences are associated with various biochemical activities, which includes regulation of gene expression ^[1]^, organization of chromosome architecture, signals controlling ^[2]^, and epigenetic inheritance ^[3]^. Amongst, these non-coding RNAs, the micro RNAs (miRNAs) present a curious case. The human genome encodes over 3000 ^[4]^ known miRNAs, of which only a small proportion has origin from non-protein-coding transcripts, whereas the rest are located mostly in the introns of coding genes ^[5]^.

miRNAs, small non-coding RNAs, came into existence as a result of evolutionary conservation of non-coding nucleotide sequences. Of all the major gene regulatory factors, the role of miRNAs in regulating gene expression has been studied extensively in last decade ^[5]^, bringing about insights of their role in regulating several cellular functions, growth, signaling, development, differentiation, proliferation, metabolism, apoptosis, and damage response ^[6-8]^. They seem to behave in a manner where they can temporarily ‘pause’ the expression of genes, thereby ensuring a temporal regulation over their expression. An overview of this regulatory behavior of the miRNAs has been schematically shown in Figure 1.

**Figure 1.**
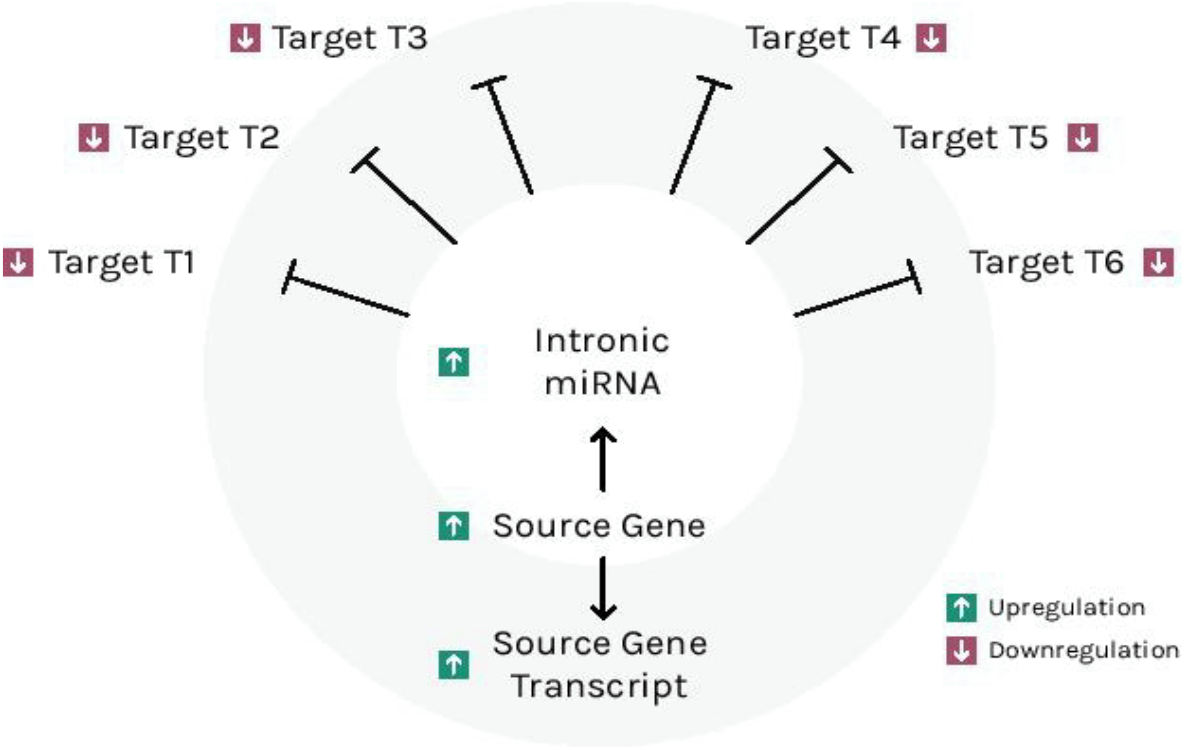
Schematic diagram showing miRNA mediated regulation of target transcripts.

With nearly 2700 experimentally validated miRNAs discovered in humans till date, many of them are shown to be associated with common human disorders. In recent years, evidences associating miRNA expressions with many diseases including immune function disorders, cardiovascular diseases, neurological diseases, viral diseases, geriatrics diseases, and cancer have been found ^[7][9-11].^ Thereby the elucidation of microRNA-disease associations is crucial for better understanding of molecular mechanisms of diseases as well as the potential for miRNA as next generation tools for diagnostic and therapeutic purposes. A large number of studies have been reported involving miRNA mediated regulation in *in-vitro* and *in-vivo* contexts that have greatly improved our understanding of miRNA–mRNA interactions ^[12]^. However, we believe that there still exists a gap in this interpretation since it does not address the highly spatiotemporal nature of these interactions in a cellular context. Furthermore, we propose that incorporating a combination of various parameters, such as thermodynamics, chemical kinetics, and co-localization, to this network would aid our understanding of these highly competitive interactors.

In order to achieve this, we have created a physicochemical scoring framework of miRNA– mRNA interaction for likelihood estimation of these interactions. Using statistical and network analysis methods we attempt to understand the regulation of gene expression by miRNAs in *Homo sapiens* at a post-transcriptional level. This analysis was used to rank these miRNAs and genes expressed within a tissue for their significance towards modulating crucial cellular functions. In the present study, we explain the details of data and formats used, and the formation of the network for tissue specific interactions. The methods used for scoring the interactions and estimating the cut off value to provide an effective network consisting of highly significant interactions have also been classified and discussed. We have also integrated several heterogeneous biological datasets to create an open-access web-server.

## RESULTS

### miRNA intron mapping and host gene identification

Since miRNAs are generally transcribed along with their host genes, the production kinetics of miRNAs was fixed to that of their host genes. Thereby, for filtering out non-intronic miRNAs from the miRNA set obtained from miRTarBase, genome coordinates of miRNAs from miRBase and intronic coordinates from UCSC Genome were overlapped. This gave us a subset of 1710 miRNAs that are intronic. For these intronic miRNAs, their host genes were secured by overlapping genome coordinates of miRNAs and coordinates of the genes from Ensembl Dataset. This process has been shown in Figure 2. This process gave us 1624 intronic miRNAs and their host genes whose expression data was retrieved from experimentally validated RNA Seq dataset from Expression Atlas ^[16-18]^. These miRNAs, their corresponding host genes, and their target genes accounted for a total of 14554 genes from the initial gene set.

**Figure 2.**
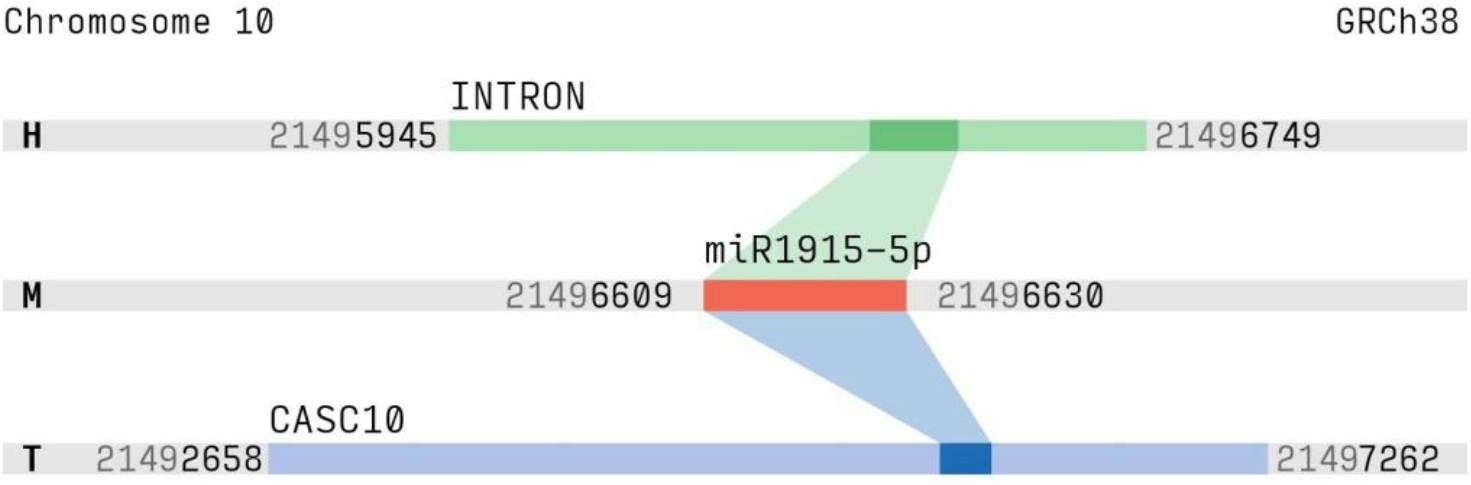
Genome Coordinate Matching of miRNA transcript (M) with host gene (H) and target gene (T) transcripts.

### Implementation

#### metaRNA

We implemented metaRNA, an open source package for computation of the thermodynamic affinity between two given RNA sequences. The package utilises the miRanda ^[22]^ algorithm and uses Vienna RNA ^[23]^ for the underlying calculations. Further information is available in Appendix 1.

#### Placet

For visualisation of large scale dataset, we developed Placet, an open source platform built using current web technologies. It uses React, D3, and eCharts for drawing content using the HTML5 Canvas API. More information is available in Appendix 2.

### Regulation Potential

We introduce the term ‘Regulation Potential’ for an miRNA–mRNA interaction as an estimate of likelihood of the interaction to hold in a given environment. A low regulation potential indicates that the interaction is less likely to hold in a given tissue, and vice-versa. It is important to note that the Regulation Potential is defined as a relative value, and hence it should not be taken as absolute.

### Tissue Specific Interaction Ranking

Many mathematical models have been proposed and are available for evaluating the competition between miRNAs. The effect of competition on regulation of expression of their targets has also been studied in specific contexts. However, a holistic view of competition among miRNAs to regulate their target genes, when taken in a given cellular context, has not been much explored. Thereby, we aim to model an effective miRNA–mRNA map depicting tissue specific analysis of interactions. The effective regulation were obtained by scoring the interactions based on thermodynamic affinity, chemical kinetics, concentration levels and network parameters like degree centrality of nodes. Since we are interested in intronic miRNAs analysis, the expression of the miRNA was taken to be equal to that of its host gene. Finally, the following expression was calculated for each interaction pairs to obtain the ranks:

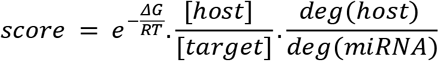

It was observed that the Relative Concentration of miRNAs and target genes, generally act as deciding factor towards the competitive regulation effects of miRNAs, while thermodynamic affinity acts as tie-breaker.

### Impact on Cellular Processes and Molecular Functions

Having obtained a Tissue Specific Interaction Ranking, we further used ontologies from PANTHER database to find the relative impact of miRNA mediated regulation over various Cellular Processes. Through the ranking of the ontology, we expect to derive a holistic overview of the functions the miRNAs play inside a cell.

The statistical overview of the ranked interactions for Liver has been presented in Figure 3. It is evident from Figure 3(a) that Apoptosis, Development, Biological Adhesion, and Immune System Processes are dominant. Whereas, when classified based on Molecular Activities, Transport, and Translation Regulation were found to be dominant, as seen in Figure 3(b). Similar observations were drawn from identical analysis of other 52 tissues from GTEx human tissue samples ^[17]^, and later validated on 32 human tissue samples from a different RNA-seq experiment (E-MTAB-2836^[18]^), details of which are available in Appendix 3.

**Figure 3.**
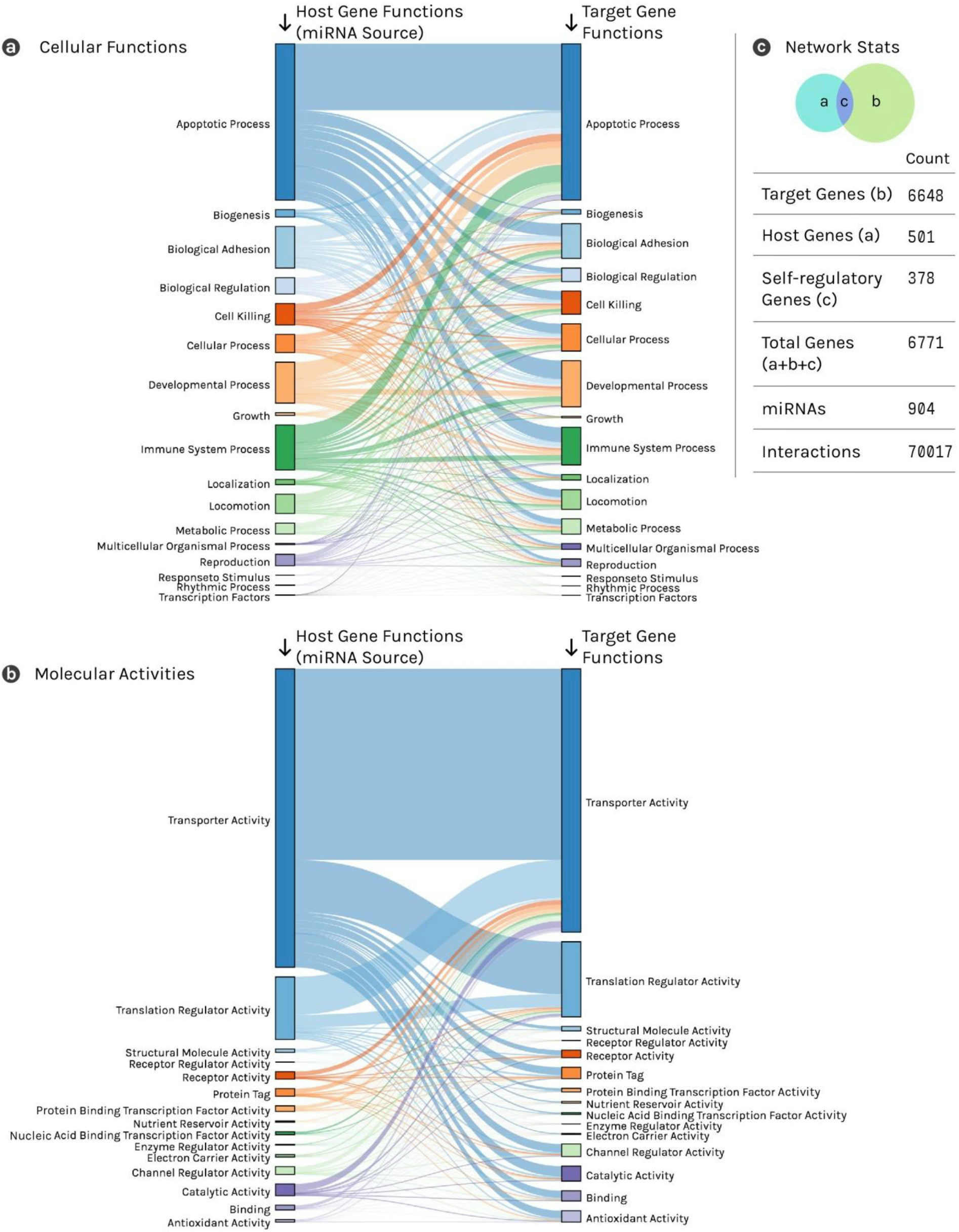
Cellular Functions and Molecular Activities in a Liver Tissue. (a) The genes involved in Apoptotic Processes, Developmental Processes, and Immune System Processes can be seen as highly co-regulatory processes, with the Apoptotic Process also being highly selfregulatory. (b) A strong self-regulatory behaviour can be seen for genes involved in Transporter Activities. Interestingly, both Transporter Activities, as well as Translation Regulation Activities are seen to be co-regulatory. (c) In the miRNA–gene regulatory network, more than three quarter of the Genes are found to be self-regulatory. A total of 501 genes account for 904 miRNAs in the regulatory network, accounting for around 70,000 interactions.

### miRNA Interaction with mRNA (miRiam) Web Server

The resultant dataset of Tissue Specific Interaction Ranking, and Ontology Ranks for each tissues are made available through the miRiam web-server. The user interface and use cases of miRiam is shown in Figure 4. Source code for the server, and databases are under an open source license. Refer to Appendix 4 for further information.

**Figure 4.**
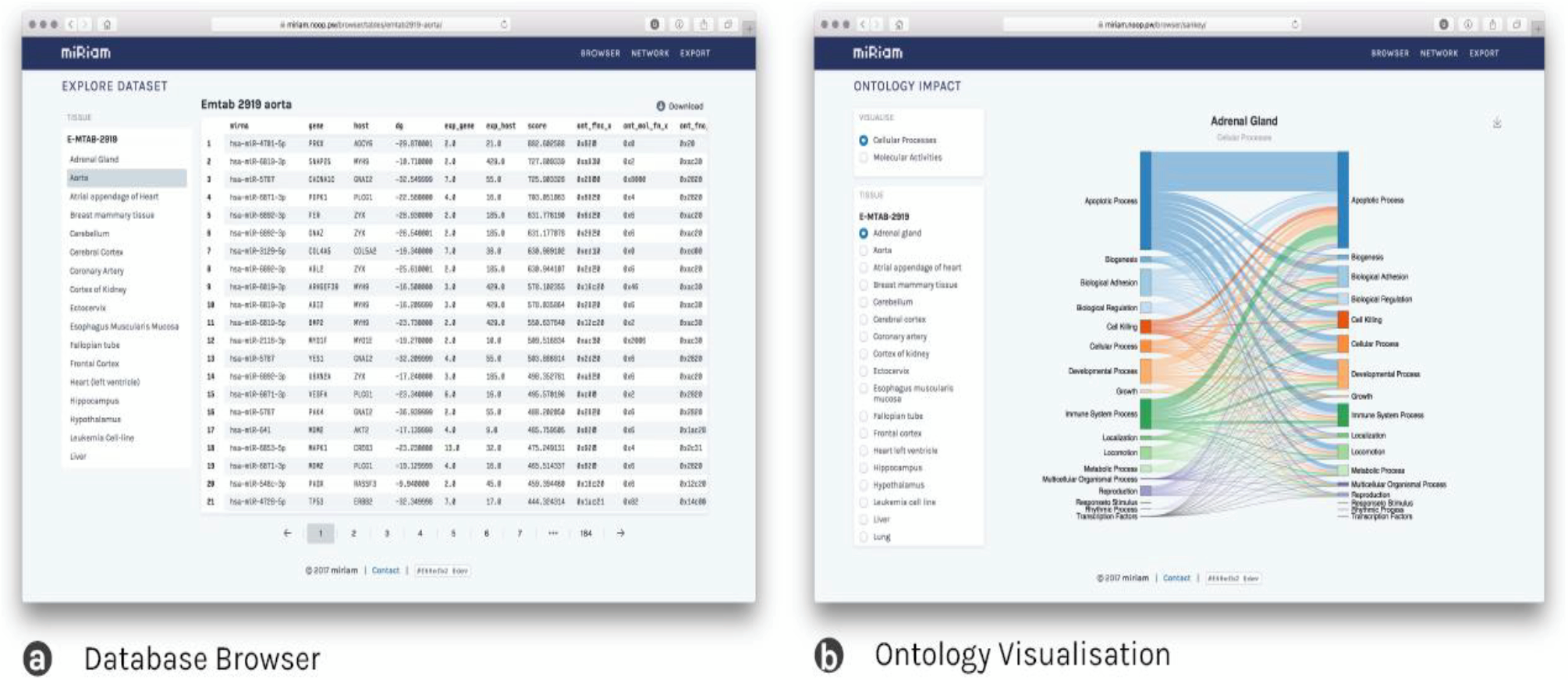
Tissue specific data exploration features as made available through miRiam. (a) Database Browser gives access to the complete raw scores, expression values, ontologies, and miRNA–gene targets. This can be exported for further usage using the provided download button. (b) Ontology visualisation gives access to interaction diagrams of Molecular Activities, and Cellular Processes for each tissue.

## DISCUSSION

### Role of miRNA as Temporal Regulators

Major components of the cellular proteins inside the cell are Ribosomal Proteins, Chaperones, and proteins involved in Glycolysis ^[24]^. Depending upon the developmental stage, the metabolic requirements of a cell vary. To ensure that the required metabolites are available within a cell, and to avoid overcrowding of nonessential metabolites, a temporal regulation of metabolite production and transport is crucial. This can be achieved through a stringent regulation of Transport Activities in the cell to selectively permit the movement of micro and macro molecules. It is important to note that this temporal regulation should also be easily reversible, should the need for specific metabolites arise.

For an optimum usage of the cell volume, it is also vital that the unnecessary molecules within the cellular system are either disposed off, or degraded for reuse. Hence degradation and waste removal processes are also essential for the longevity of the cells. Furthermore, these processes are also required to be temporal.

Gene Expression Regulation through Transcription Factors, mRNA Stability and Rate of Degradation, and DNA Modification give rise to a permanent change in cellular gene expression levels, which can be visualised as ‘start’ and ‘stop’ functions. While necessary for cell differentiation, such regulatory activities are not easily reversible, making them unsuitable as temporal regulators. By binding to a mRNA transcript, a miRNA ‘pauses’ its expression by forming a mRNA–miRNA complex, thereby controlling specific protein production in a temporal fashion. This complex formation, however, is easily reversible, thus indicating that the miRNAs act as temporary regulators of gene expression.

### Role of miRNA in management of Protein Crowding inside the cell

Our analysis reveal that the intronic miRNAs largely regulate the translation of abundantly produced proteins that are involved in major metabolic processes, transport activities, translational machinery, and catalytic activity. Since these proteins are required in a temporal fashion throughout a cell’s lifespan, their production cannot be completely shut down. However, a continuous production of these proteins is also not desirable, as that would cause crowding inside the cell, hindering the transitory movement of essential molecules.

Hence, a temporal regulatory mechanism is immensely important to stow away mRNAs that are not required to be translated immediately, but would be required later. In this way, such regulation prevents over-crowding of the cell by unnecessary proteins. miRNA mediated gene expression regulation, as a cascade of temporal regulator, has an important role in ensuring the optimal usage of space within the cell. This has also been substantiated by the experimental evidences that show intronic miRNAs’ involvement in apoptosis, protein accumulation and maintenance of cell size ^[25-31]^.

## MATERIAL AND METHODS

### Datasets

#### miRTarBase

A collection of miRNAs and their target genes in *homo sapiens* was obtained from miRTarBase ^[13]^. This was utilised to construct an overview network consisting of 2649 miRNAs and 14894 target genes.

#### miRBase

A comprehensive list of chromosome coordinates of miRNAs were acquired from the miRBase database ^[4]^. In addition to this, we also obtained the mature RNA sequences for these miRNAs.

#### UCSC Genome Browser

We acquired chromosome intron coordinates from University of California Santa Cruz Genome Browser database ^[14]^.

#### Ensembl

CDNA sequences of the genes in our network were downloaded using Ensembl ^[15]^ via REST API. The chromosome coordinates of these genes were also obtained.

#### Expression Atlas

We obtained gene expression data for following RNA-Seq experiments from Expression Atlas ^[16]^:

- RNA-seq from 53 human tissue samples from the Genotype-Tissue Expression (GTEx) Project (E-MTAB-2919 ^[17]^).
- RNA-seq of coding RNA from tissue samples of 122 human individuals representing 32 different tissues (E-MTAB-2836 ^[18]^).

#### CPDB

Genes annotations based on their participation in particular biochemical pathways were obtained from ConsensusPathDB ^[19]^. These annotations were used to group together genes which participate in a particular pathway.

#### NCBI GenBank

Gene description, functions, synonyms, and other metadata for the genes were downloaded from NCBI GenBank ^[20]^.

#### PANTHER

Functional classification of the genes were obtained from PANTHER ^[21]^. These functional classes tag genes with their involvement in cellular processes, for example ‘Apoptotic Process’, ‘Reproduction’, etc., and with their involvement in molecular activities, such as ‘Antioxidant Activity’, ‘Transporter Activity’, etc.

### Equilibrium Constant as a characteristic for significant interaction

The equilibrium constant is the value of the reaction coefficient when the reaction has reached equilibrium and therefore, is a measure of the stability attained by the interaction. Thermodynamic affinity for each interaction was calculated using metaRNA. Further, the following expression is used to calculate the equilibrium constants, *k*_eq_:

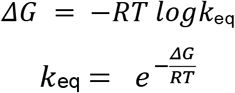

### Expression as a characteristic for significant interaction

The kinetics of a reaction govern the rate and direction in which the reaction proceeds and hence, is an important factor in determining the interaction scores. Therefore, the ratio of expression level of miRNA to target gene plays a crucial role, if the expression of miRNA is greater than that of the target gene, the regulation potential of the miRNA is higher. This means that the miRNA can function as a pause function and can therefore regulate the expression of the gene completely. Alternatively, if the miRNA is expressed in lesser quantity than its target gene, the miRNA would only be fine tuning the expression and the regulation potential would be much lower.

### Functional Classes and Pathways as a characteristic for significant interaction

Ontologies for Functional Classes, Molecular Activities, and Related Pathways of genes were obtained from PANTHER Database. These ontologies were used as ‘Concurrence Parameter’. The concurrence parameter is an indicative quantification denoting the likelihood of the miRNA, gene to be co-expressed and co-located. An interaction is given a higher concurrence score if its components share a common function, molecular activity, or occur in same pathways.

### Network Parameters as a characteristic for significant interaction

Network Parameters like degree centrality measures are crucial in determining the significance of an interaction. If a miRNA targets a large number of genes, its ability to regulate a specific target would naturally reduce since the same quantity of the miRNA would be responsible to regulate a large number of targets. Hence, the role played by this miRNA for a particular interaction would be lesser as compared to an miRNA which targets lesser number of genes. In contrast to this, if a particular gene has a large number of miRNAs targeting it, the cumulative regulation effect through multiple miRNAs increases.

## DATA AVAILABILITY

### METARNA

Released under a permissive open source license (MIT), metaRNA is available for download from Python Package Index (PyPi) https://pypi.python.org/pypi/metarna. The source-code is also available at https://github.com/prashnts/metaRNA.

### PLACET

At its core, Placet uses echarts https://github.com/ecomfe/echarts for visualisation using HTML5 Canvas API. D3 https://d3js.org/ is used for utility functions such as scales and colour pallets. Finally React https://reactjs.org/ is used for a consistent two-way data binding that controls the whole user interface.

### miRiam WEB SERVER

The dataset made available through miRiam web server is accessible from https://miriam.noop.pw. Source code for this is also available from https://github.com/prashnts/miriam-explore. The webserver is written using Django, a high performance web framework in Python, and utilises Placet for implementation of the user-interface.

## ACKNOWLEDGEMENT

Authors thank Dr. Vinod Scaria, and Dr. Beena Pillai for their valuable inputs.

