## Supplementary Materials for "Intronic miRNA Mediated Gene Expression Regulation Controls Protein Crowding Inside the Cell"

**SUPPLEMENTARY DATA**

### APPENDIX 1: METARNA

metaRNA finds potential target sites for the microRNAs in genomic sequences. It is built using miRanda [22], an algorithm for detection and ranking of the targets of microRNAs. metaRNA is written in Python, and C programming languages.

Released under a permissive open source license (MIT), metaRNA is available for download from Python Package Index (PyPi) https://pypi.python.org/pypi/metarna.

The source–code is also available at https://github.com/prashnts/metaRNA. The miRanda [22] algorithm works in two phases. In phase one, the potential target sites are reported based on query microRNA and reference (CDNA) sequence. These targets are scored and the high scoring alignments are then used in second phase, where the folding routines of RNAlib [23] library are utilised to calculate the minimum free energy of the resulting combinations.

### APPENDIX 2: PLACET

At its core, Placet uses echarts https://github.com/ecomfe/echarts for visualisation using HTML5 Canvas API. D3 https://d3js.org/ is used for utility functions such as scales and colour pallets.
Finally React https://reactjs.org/ is used for a consistent two-way data binding that controls the whole user interface.

### APPENDIX 3: MOLECULAR PROCESSES VISUALISATION

We have given a weighted bipartite graph $G$, represented using an edgelist data structure. For each node $n_i$ in $G$, we also have a set $M=\{x_i, m\}$ which represents $m$ decimal value attributes as annotations of $n_i$. We apply a transformation to $G$, to obtain a weighted graph $G'$ where:

1. There are 2$m$ nodes: $\{x_{i\text{-source}}, m\}\ U\ \{x_{i\text{-sink}}\}$.
2. An edge between $x_{i\text{-source}}$, and $x_{j\text{-sink}}$ is defined if, in $G$, a node having a finite value annotation for $x_i$, and links another node with finite annotation for $x_j$ . The weight of this edge is taken as sum of all edge weights in $G$ where such condition holds true.

This transformation gives us data required to render the Sankey diagram using Placet, as shown in Figure 3.

**APPENDIX 4: miRiam WEB SERVER**

The dataset made available through miRiam web server is accessible from https://miriam.noop.pw. Source code for this is also available from https://github.com/prashnts/miriam-explore. The webserver is written using Django, a high performance web framework in Python, and utilises Placet for implementation of the user-interface.
